# Interactions in CSF1-driven Tenosynovial Giant Cell Tumors

**DOI:** 10.1101/2022.06.01.494428

**Authors:** David G.P. van IJzendoorn, Magdalena Matusiak, Gregory W. Charville, Geert Spierenburg, Sushama Varma, Deana R.C. Colburg, Michiel A.J. van de Sande, Kirsten van Langevelde, David G. Mohler, Kristen N. Ganjoo, Nam Q. Bui, Raffi S. Avedian, Judith V.M.G. Bovée, Robert Steffner, Robert B. West, Matt van de Rijn

## Abstract

The majority of cells in Tenosynovial Giant Cell Tumor (TGCT) are macrophages responding to CSF1 that is overproduced by a small number of neoplastic cells with a chromosomal translocation involving the *CSF1* gene. Treatment with inhibitors of the CSF1 pathway has been clinically effective. An autocrine loop was postulated where the neoplastic cells are stimulated through the CSF1 receptor (CSF1R) expressed on their surface. Here we show that the neoplastic cells themselves do not express CSF1R and therefore may be unaffected by current therapies. We identified a new marker for synoviocytes, GFPT2, that highlights the tumor cells in TCGT and is associated with activation of the YAP1/TAZ pathway. The neoplastic cells in TGCT are highly similar non-neoplastic synoviocytes. Finally, we provide molecular support for the osteoclast-like features of the giant cells in TGCT that correlate with the destructive effects of TGCT on bone.

## Introduction

Tenosynovial giant cell tumor (TGCT) occurs in the synovial tissue within joints or tendon sheaths. While they do not metastasize, they can destroy cartilage and bone and can lead to debilitating effects on joint function. Moreover, TGCT’s often aggressively recur after surgery (1). Our group previously described a small subpopulation of cells within the tumor that carry a translocation involving CSF1. These neoplastic cells express high levels of CSF1 that is known to attract and induce proliferation of the monocytes expressing the receptor for CSF1 (CSF1R) on their surface (2). In this way the neoplastic cells have a “landscaping effect” on their microenvironment that results in the accumulation of large numbers of the non-neoplastic macrophages that form the majority of cells in the tumor (3, 4). Initially, we identified COL6A3 as the fusion partner for *CSF1* but others have subsequently identified a range of other translocation partners for *CSF1* in TGCT (5, 6). The findings suggested that the chemoattractant and proliferative effect of CSF1 on bystander macrophages could be inhibited by drugs that interfered with the CSF1 pathway. Studies using monoclonal antibodies and small molecule inhibitors for the CSF1 pathway have shown a significant clinical effect (6–13) and a decrease in tumor size as seen by an imaging measurement specifically designed for TGCT (14). It was hypothesized that the CSF1-expressing neoplastic cells might be activated through an autocrine loop by CSF1R expressed on their surface (3). However, the effect that drugs targeting the CSF1 pathway have on the neoplastic cells has not been studied. Understanding whether the anti-CSF1R therapies also target the neoplastic compartment will help rationalize the duration of the therapy administration, and asses the risk of the recurrence. Here we used scRNA sequencing combined with a long-read mRNA sequencing of 3 TCGT tumors and analysis of patient samples before and after anti-CSF1R therapy. We established a new FFPE-compatible IHC marker for detecting the neoplastic cells in the TGCT. We show that inhibitors of the CSF1R are unlikely to directly affect growth of the neoplastic cells because CSF1R in the TGCT is mainly expressed on the myeloid compartment. We show that CSF1R targeting in TGCT causes a relative increase in neoplastic cell frequency in the tumor. We identify 2 pathways that suggest new candidates for treatment of these cells. In addition, we further provide insight into the relationship between the neoplastic cells and synoviocytes and into the features that define the giant cells in this tumor.

## Results

### Single cell RNA-seq and long-read RNA-seq identify the neoplastic cells in Tenosynovial Giant Cell Tumors

To define the cellular interactions in TGCT, fresh tumor material was collected from three TGCT patients (TGCT1-3). The presence of the characteristic translocations involving *CSF1* was confirmed on FFPE material by FISH (sup fig 1) in each case. Five 10x barcoded libraries were generated from the three cases and used for single cell RNA sequencing resulting in a total of 11,430 cells for analysis. The cells clustered in 10 distinct groups by Umap analysis (fig 1a). To determine the exact site of the translocation within the *CSF1* gene and the fusion partners, full length mRNA was profiled using long-read RNA-Seq (PacBio Iso-Seq) (fig 1b). The known fusion sequence from each case was used as a custom reference genome and scRNA reads were aligned to these reference genomes to identify the neoplastic cells that contain the CSF1 translocation. In total, 79 cells were identified that contained split reads (253 split reads in total) involving CSF1. Cells with CSF1 translocations were identified for *CSF1::FN1* in 11 cells (TGCT1). In TGCT1, *CSF1::PDPN* was detected in 22 cells (TGCT2). The fusion breakpoint is halfway exon 6 of *CSF1* and connects to exon 2 of *PDPN*. The transmembrane domain for CSF1 is lost but the fusion partner (PDPN) donates an in-frame transmembrane domain. In TGCT3, *CSF1::EBF1* was detected in 46 cells (TGCT3). The 79 cells with confirmed fusion were restricted to two clusters in the Umap analysis that also showed high expression of CSF1 and other unique markers (fig 1a, insert, sup fig. 2). The remaining cell clusters were identified based on expression of canonical marker genes (sup fig 2) resulting in distinct cell populations consisting of macrophages, proliferating macrophages, giant cells, T-lymphocytes, fibroblasts, smooth muscle cells, pericytes, endothelial cells and the two clusters with neoplastic cells (fig 1c). Concordant with previous findings, the cells that carry the CSF1 translocation as found by long-read RNA-seq represent a minority of cells withing the tumor mass. These cells map to 2 clusters that together make up 10% of the total cells on average. It cannot be fully excluded that the two neoplastic cell clusters also contain normal cells. As reported previously, the largest cell population of cells within TGCT consist of macrophages (35%, fig 1d). Cell cycle state analysis with Seurat showed that the proliferating cell cluster in this analysis consisted exclusively of macrophages (sup fig 2, sup fig 3a) and was absent from the tumor cells as confirmed by Ki67 immunofluorescence (sup fig 3b).

**FIG 1.**
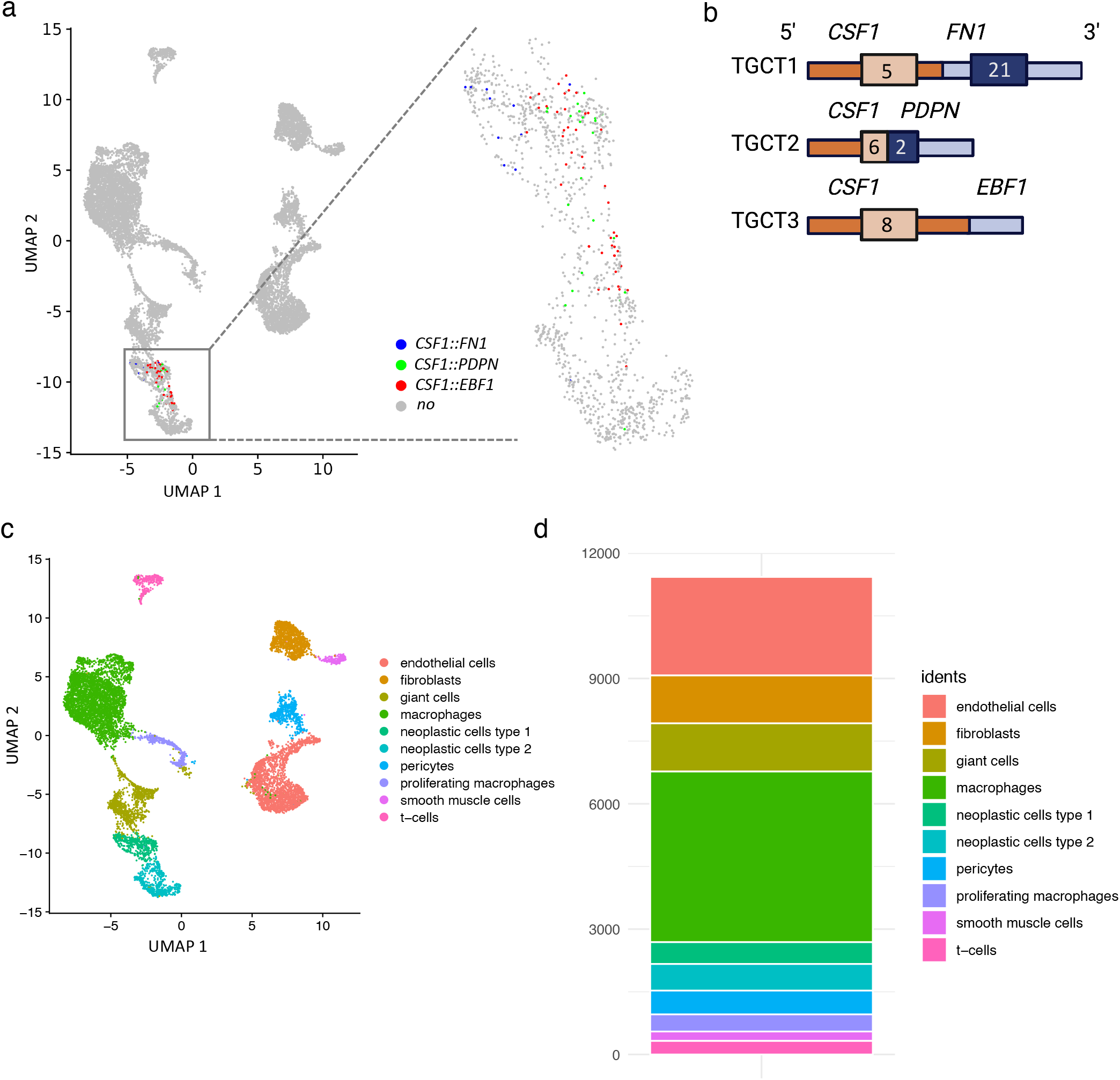
Overview of Tenosynovial Giant Cell Tumors single cell RNA-seq and long-read mRNA sequencing data. (a) UMAP plot with scRNA-seq data from three TGCT patients. The presence of gene fusions involving the *CSF1* gene is plotted. The cells harboring fusions are located in two adjacent clusters (box), a blowout of these clusters is shown on the right. (b) The three fusions involving *CSF1* identified with long-read mRNA sequencing are shown schematically. (c) Identified scRNA-seq clusters were named according to their canonical markers, the presence of cells with fusions and s score. (d) Bar graph shows the number of cells identified in each of the cell clusters in the scRNA-seq data.

### TGCT contains two populations of neoplastic cells that correlate with two previously identified subtypes of synoviocytes

Previous studies already suggested that the neoplastic cells in TGCT are derived from synoviocytes but this was based on expression of relatively few markers (clusterin and desmin) by immunohistochemistry without confirmation of the neoplastic nature of the cells (15, 16). To further study the two neoplastic cell clusters, we compared the neoplastic cells to a publicly available scRNA dataset on synovial tissue samples from rheumatoid arthritis (RA) patients by Stephenson, et al. (17). Despite a difference in technologies used, there was a significant overlap between macrophages, endothelial cells, and smooth muscle cells from the 2 studies after integration using Seurat (data not shown). In addition, the two synoviocyte subpopulations found in the RA data clustered closely with the neoplastic cell clusters identified in the TGCT cases (fig 2a). To further compare the synoviocytes from both datasets a SVM classifier was trained to characterize the two types of RA synoviocytes. With this classifier it was found that the majority of cells within each of the two neoplastic cell types clustered together in either of the two synoviocyte clusters from Stephenson et al (fig 2b).

**FIG 2.**
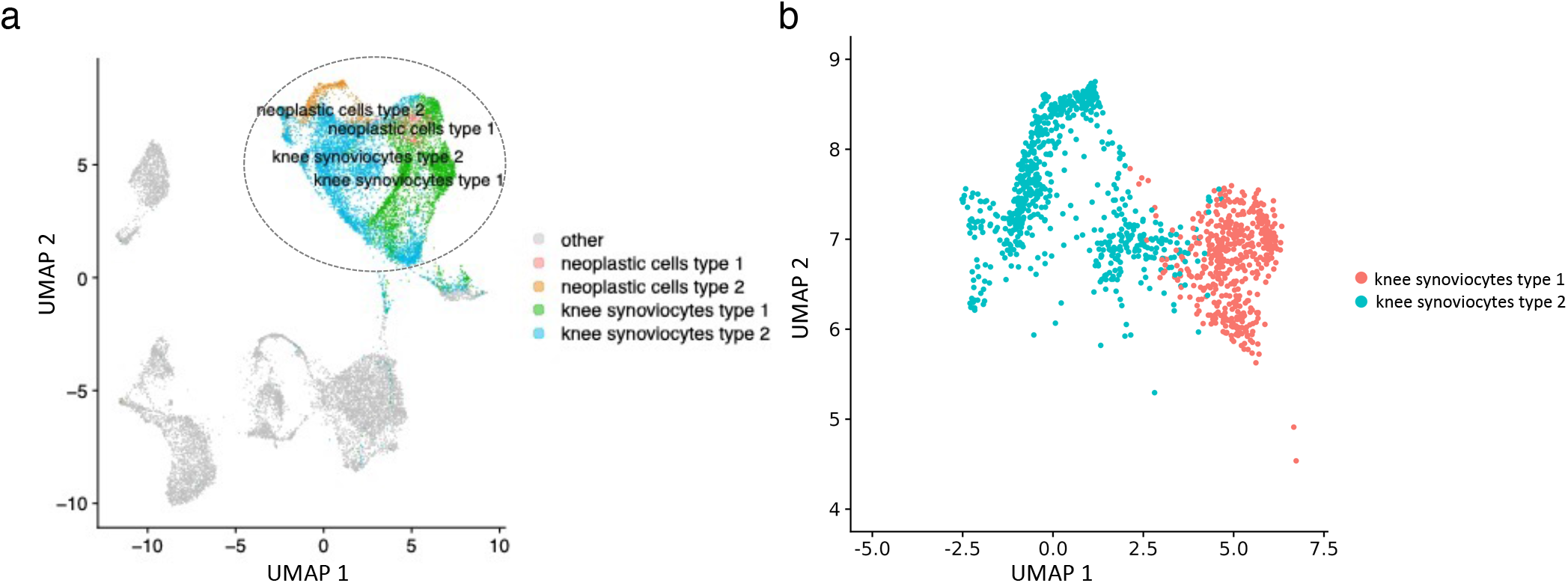
ScRNA-seq data from the TGCT cases was compared to published synoviocyte scRNA-seq data from RA patients (17). The two subtypes of RA synoviocytes correspond to the two clusters of neoplastic cells that were identified in TGCT samples. (a) scRNA-seq data from synoviocytes identified in RA was merged with the TGCT scRNA-seq data, showing that the neoplastic cells and synoviocytes cluster close together (dashed circle). (b) A SVM classifier was trained on synoviocytes from RA dataset and applied to the neoplastic cells in TGCT. The neoplastic cells type 1 and 2 correspond to the two subtypes of synoviocytes found in RA.

A gene expression analysis showed that the cells in the 2 neoplastic clusters uniquely expressed GFPT2 and high levels of CSF1. Comparison between the 2 neoplastic clusters showed that type 1 had higher expression of *CD68, CD55, CLU* and *TREM1* while neoplastic cells in cluster 2 were characterized by high levels of expression of *COL6A3, POSTN, FGF7, PODN, RARRES1, CTHRC1, SLIT3, LAMB1* and *ADAM12* (sup fig 4). Combined with the results from long mRNA profiling, our data shows that the *CSF1* translocation for each of the 3 patients occurs in both synoviocyte subtypes and stresses the high similarity between the neoplastic cells in TGCT and normal synoviocytes.

### GFPT2 is a new marker for neoplastic cells in TGCT and is associated with activation of the HIPPO signaling pathway

*GFPT2* mRNA was found to be highly expressed in both neoplastic cell types (fig 3a) with the highest level of expression in type 1 neoplastic cells. In contrast to two other genes *CLU* and *PDPN* that were previously reported in TGCT (15), GFPT2 is specific for the neoplastic cells in TGCT and, unlike CLU or PDPN, is not expressed in giant cells (fig 3a, sup fig 5a). Others have reported expression of desmin in TGCT (15, 16) but no desmin mRNA expression was found in the neoplastic cell clusters (sup fig 5a) and IHC against desmin was negative (data not shown). The lack of desmin IHC staining might be explained by differences in antibodies used in different laboratories.

**FIG 3.**
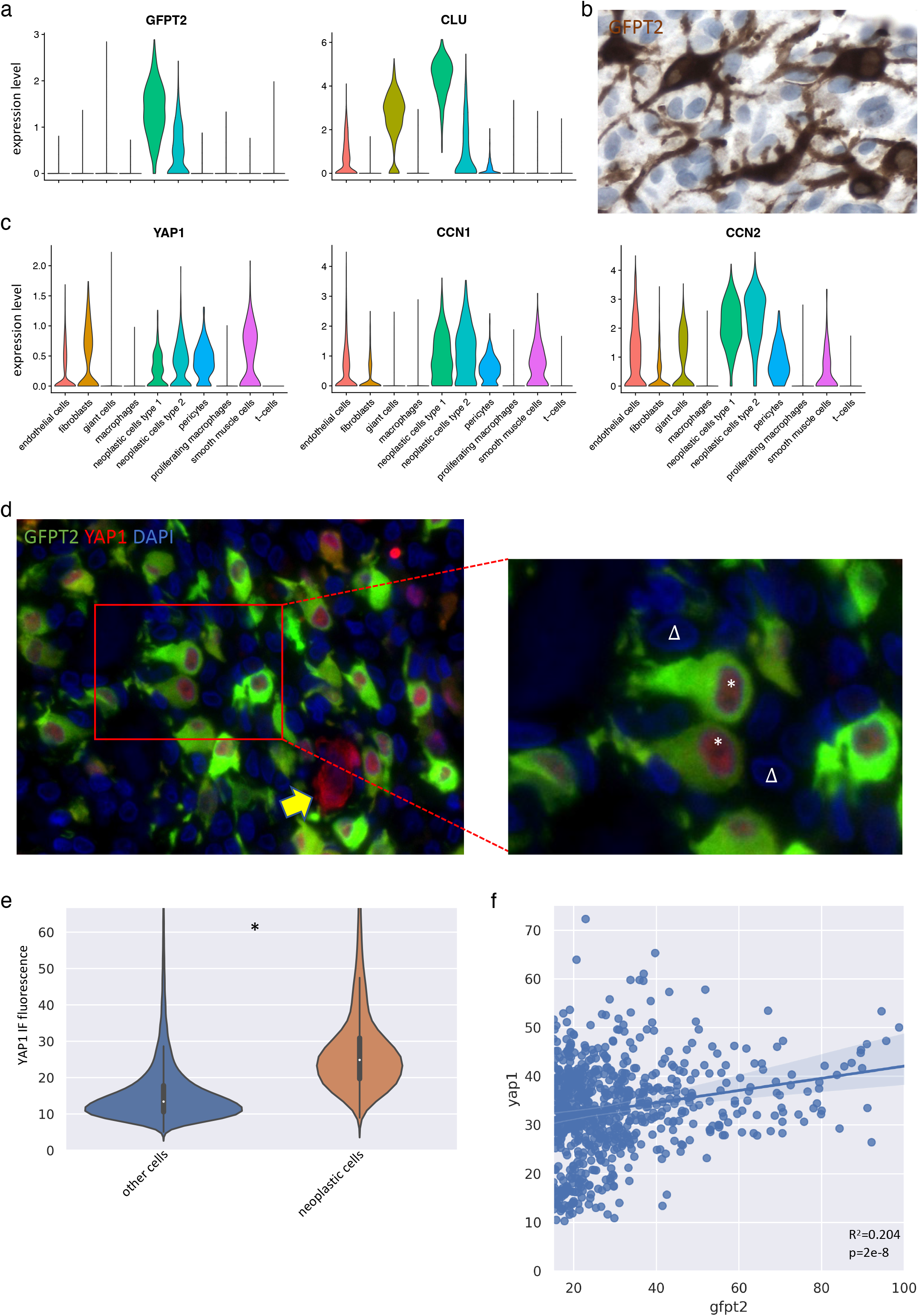
GFPT2 is a specific marker for TGCT neoplastic cells. GFPT2 highlights the dendritic nature of the neoplastic cells and its high expression is associated with activation of the HIPPO signaling pathway. (a) ScRNA-seq gene expression levels for *GFPT2* and *CLU* in all cell types found in TGCT. *GFPT2* is specific to the two neoplastic cell clusters with higher expression in neoplastic cell cluster 1. *CLU* is highly expressed in the neoplastic cells as well but is also present in giant cells and endothelial cells. (b) IHC highlights the dendritic appearance of the GFPT2 positive cells. The GFPT2+ cells are surrounded by GFPT2-cells with smaller nuclei. (c) scRNA-seq gene expression levels for *YAP1, CCN1* and *CCN2* showing expression in both neoplastic cells clusters. (d) IF stain for GFPT2 and YAP1 shows nuclear localization of YAP1 in the neoplastic GFPT2+ cells (*) in an area chosen to contain a dense concentration of neoplastic cells. The surrounding GFPT2-cells do not show nuclear expression of YAP1 (Δ). Some GFPT2 negative cells show YAP1 in the cytoplasm but not in the nucleus (yellow arrow). (e) YAP1 IF signal quantification in GFPT2- and GFPT2+ cells. GFPT2+ cells show a significantly higher (* = p<0.05) expression of nuclear YAP1. (f) Quantification of the IF signal intensity for cytoplasmic GFPT2 and nuclear YAP1 within the group of GFPT2+ cells. A significant (p = 2e-8) Pearson correlation was found between the signal intensities (R^2^ = 0.204).

IHC for GFPT2 showed that the neoplastic cells have a dendrite like morphology (fig 3b) and cells with a remarkable dendritic appearance were also noted in the single cell preparations for the scRNA-Seq libraries (sup fig 5b). GFPT2 is a rate-limiting enzyme in the hexokinase pathway but also functions as a regulator of the HIPPO signaling pathway (18). YAP1, the most important transcription factor in the HIPPO signaling pathway, is localized to the nucleus when activated. *YAP1* mRNA is expressed in the neoplastic cells cluster and downstream genes from the HIPPO signaling pathway (*CCN1* and *CCN2*) were also found to be upregulated in the neoplastic cells (fig 3c). Immunofluorescence (IF) showed nuclear localization of YAP1 protein in the GFPT2 positive cells (fig 3d, see *), indicating activation of the pathway (18). Many GFPT2-negative cells had no YAP1 expression, (fig 3d, see Δ). While GFPT2-negative cells can show *YAP1* mRNA expression (fig 3c), the majority of these cells showed cytoplasmic staining only (fig 3d, arrow) indicating non-activated YAP. Quantification of the nuclear IF signal intensity for YAP1 confirmed significantly higher intensity in the neoplastic (GFPT2-positive) cells than in non-neoplastic cells (p<0.05, fig 3e). Within the GFPT2-positive cell population there also was a significant correlation between GFPT2 and nuclear YAP1 IF signal (fig 3f, R^2^=0.204, p=2e-8).

### CSF1 stimulates macrophages and giant cells but does not affect neoplastic cells through an autocrine loop

In our previous studies we suggested that neoplastic cells in TGCT could be stimulated through secreted CSF1 via an autocrine loop (3). <u>CellphoneDB</u> was used in the current study to identify ligand-receptor interactions between the two neoplastic cell subtypes and other cell types for CSF1, IL34 and PDGFA and -B. Significant interactions were identified between neoplastic cell subtypes 1 and 2 and the macrophages and giant cells involving CSF1 and CSF1R (fig 4a). Neoplastic cell subtype 2 also interacted with the macrophage giant cell populations through IL34 signaling (fig 4a). In contrast, no interactions were found in CellphoneDB to suggest an effect of CSF1 on the neoplastic cells themselves. High expression of *CSF1* localized to both neoplastic cell subtypes (fig 4b). *IL34* was expressed in neoplastic cells in cluster 2 but not in cluster 1. *CSF1R* is highly expressed in the giant cells, macrophages and proliferating macrophages but is absent from the neoplastic cells and other cell populations (fig 4b).

**FIG 4.**
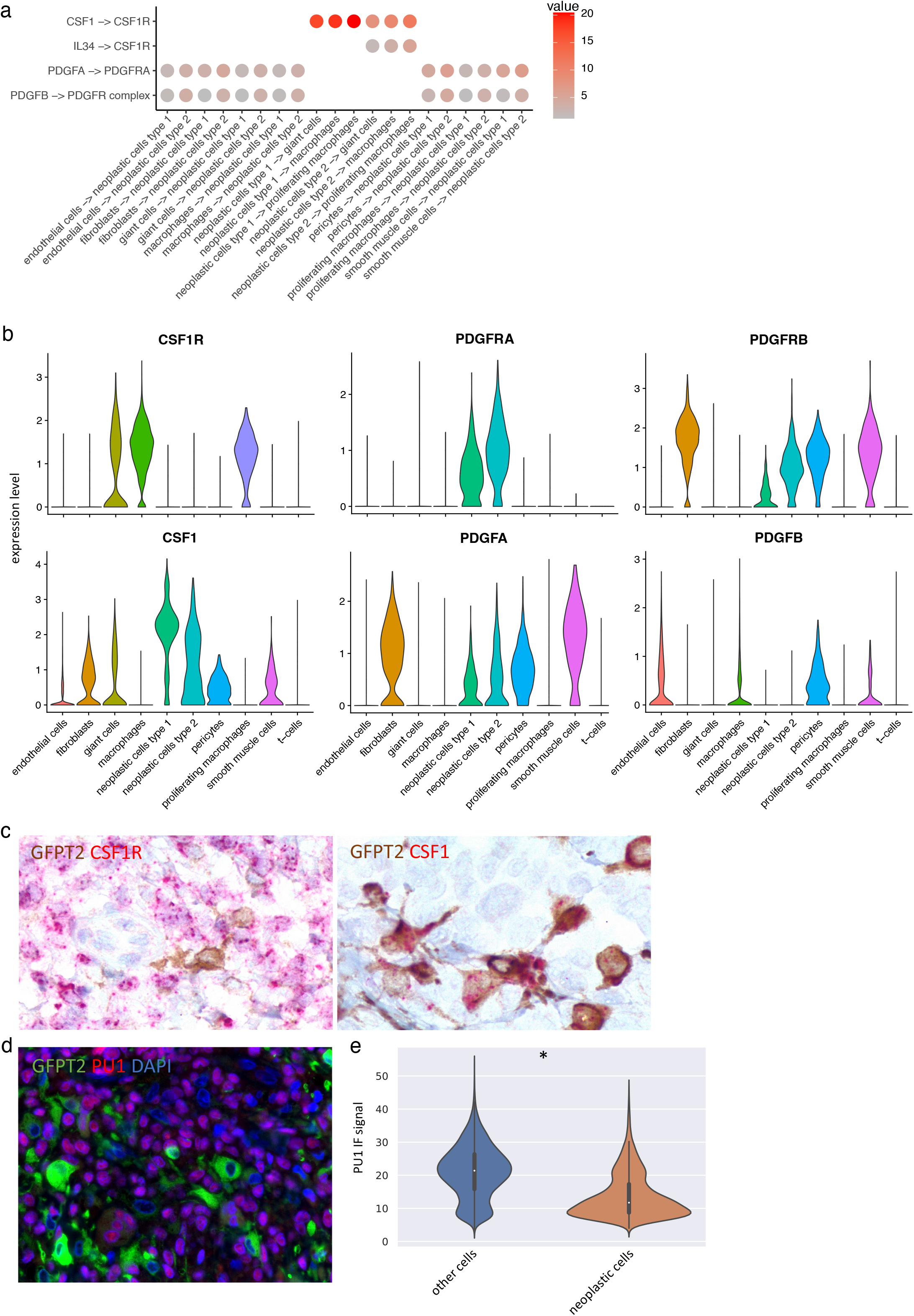
Ligand-receptor interactions for the neoplastic cells in TGCT show absence of an autocrine loop involving CSF1 signaling. (a) CellphoneDB identified interactions involving CSF1, IL34 and PDGF-A and -B between the neoplastic cell clusters and other cell subtypes. (b) scRNA-seq expression of the receptors (*CSF1R, PDGFRA* and *PDGFRB*) and ligands (*CSF1, PDGFA* and *PDGFB*). (c) Combined GFPT2 (IHC) and *CSF1R* or *CSF1* (RNA-ISH). GFPT2 and *CSF1* co-localize while the GFPT2+ cells are negative for *CSF1R* mRNA. (d) IF for GFPT2 and PU.1. GFPT2+ cells are negative for PU.1. (e) Nuclear PU.1 signal intensity was quantified for the GFPT2+ and GFPT2-cells, showing a significantly higher (* = p < 0.05) nuclear signal in the GFPT-cells.

Two additional orthogonal approaches were used to confirm that the neoplastic cells do not express detectable CSF1R. First, GFPT2 IHC combined with RNA-ISH for *CSF1R* showed that GFPT2 and *CSF1R* were not present in the same cell populations (fig 4c) validating the scRNA findings. In contrast, IHC for GFPT2 combined with RNA-ISH for *CSF1* showed co-localization of GFPT2 and *CSF1*. Second, *SPI1*, a gene in the regulatory network of the CSF1R receptor showed high expression in the macrophages and giant cells. PU.1, the protein corresponding to *SPI1*, was not present in the nucleus of the GFPT2-positive neoplastic cells but showed nuclear localization in the surrounding cells (fig 4d). Quantification of the PU.1 IF signal intensity showed significantly higher expression in the GFPT2-negative cells (fig 4e). These findings indicate the absence of an autocrine CSF1 signaling loop in the neoplastic cells and suggest that drugs that interfere with the CSF1 pathway may affect the bystander macrophages and giant cells in TGCT but not the neoplastic cells.

### Support for proliferative signals from the microenvironment to the neoplastic cells

Previous studies have shown that tyrosine kinase inhibitors such as imatinib that have a relatively broad target spectrum can inhibit TGCT. A major target of imatinib is KIT but there is no significant expression of this gene in TGCT (data not shown). Other targets of Imatinib are CSF1R, PDGFRA and –B. Interaction analysis (CellphoneDB) for PDGFA and –B indicated interactions between PDGF receptors on the neoplastic cells and other cell types in TGCT (fig 4a). ScRNA expression levels for these 4 genes validated the interaction analysis findings. The receptors for PDGFA and –B are expressed on both types of neoplastic cells with PDGFRA being exclusive for the neoplastic cells in TGCT. Other components of the microenvironment express measurable levels of PDGFA, -B or a combination of the two (fig 4b). These findings suggest a supportive effect from the non-neoplastic cells in the microenvironment on the neoplastic cells and raise the possibility for a combination therapy targeting CSF1 to inhibit macrophages and PDGFRA to inhibit the neoplastic cells.

### Clinical specimens suggest survival preference for neoplastic cells after treatment with inhibitors of the CSF1 pathway

To study the effect of CSF1R inhibition on TGCT tumors *in vivo*, material from four patients was analyzed (clinical detail in Supplementary table 1). Two patients (Patient 1, 2) had been treated with pexidartinib, a small molecule inhibitor of CSF1R. One patient (Patient 3) was treated with nilotinib, another small molecule inhibitor for CSF1R. The two patients that were treated with pexidartinib showed a reduction in tumor volume according to the Tumor Volume Score (TVS) and the TVS for the Nilotinib treated patient remained the same (fig 5a and fig 5b). For these patients samples were available prior to treatment and from variable periods after treatment (Supplementary table 1). The patient samples were stained for GFPT2, PU.1 and DAPI. A representative image shows an apparent increase in the percentage of GFPT2-positive neoplastic cells compared to PU.1-postive macrophages in the specimens after treatment with pexidartinib or Nilotinib (Fig 5c). The number of GFPT2-positive (neoplastic cells), PU.1-positive (macrophages and giant cells) and double negative (other) cells were quantified by measuring IF signal for each cell in representative 2.94mm^2^ areas. The total number of cells was used to determine the fraction of neoplastic cells, and macrophages and giant cells in each representative area. For all three cases the fraction of macrophages remained relatively stable while the fraction of neoplastic cells increased (fig 5d).

**FIG 5.**
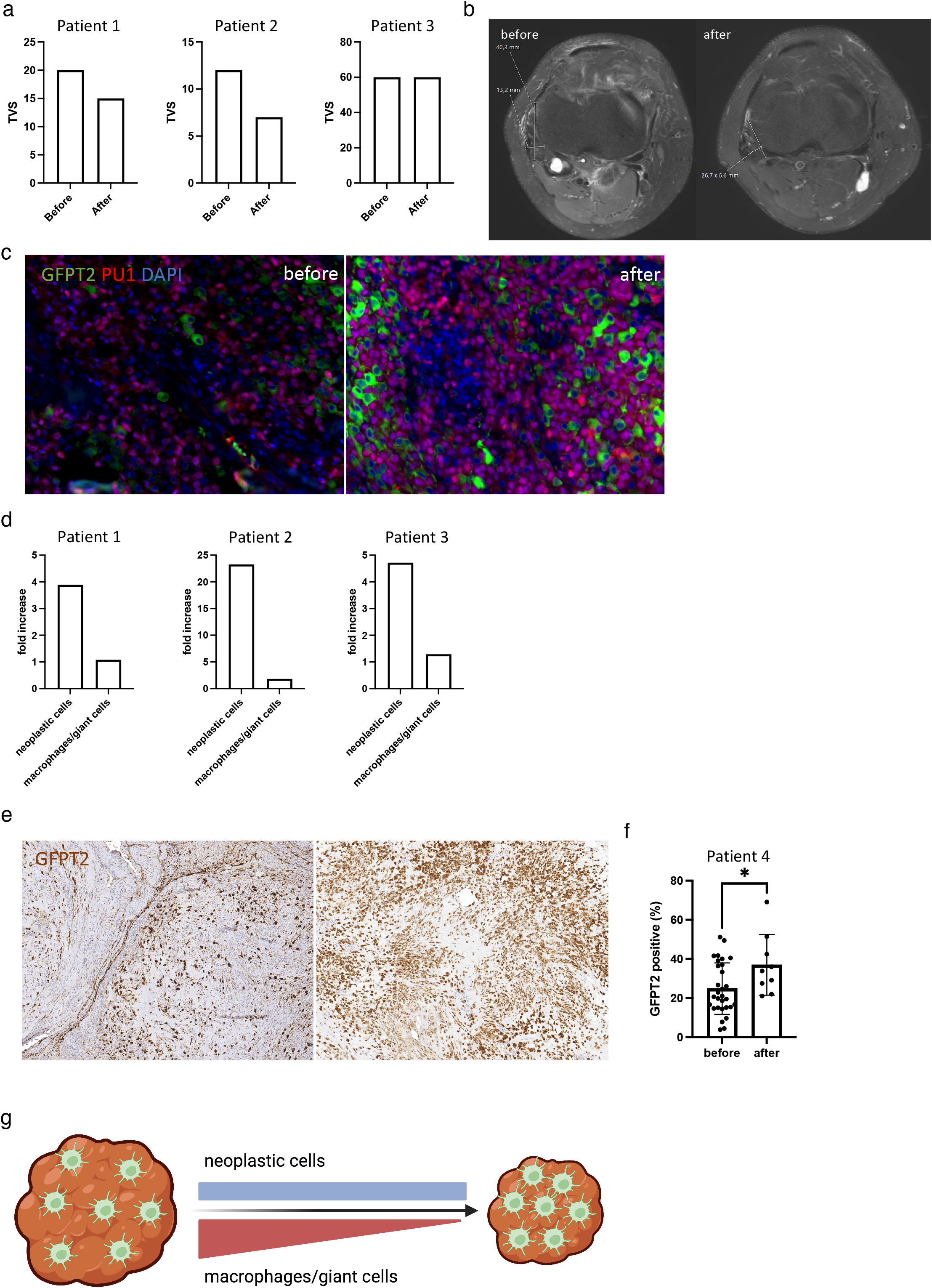
Small molecule and antibody treatments targeting the CSF1R pathway led to a radiological decrease in tumor volume. This treatment however does not target the neoplastic cells. In 4 TGCT patients treated with three different CSF1R inhibitors an increase in number of neoplastic cells per microscopic field is shown. (a) Patients 1 and 2 were treated with pexidartinib and patient 3 was treated with Nilotinib. Tumor Volume Score (TVS) decreased in patients 1 and 2 and remained stable in patient 3 during the treatment. (b) MRI image from patient 2 before (left) and after (right) systemic pexidartinib treatment. (c) IF for GFPT2 and PU.1 was performed on patients 1-3. Left panel shows a representative image before pexidartinib treatment and right panel shows the same patient after pexidartinib treatment. (d) The fraction of GFPT2-positive and PU.1-positive cells was determined before and after treatment in a total area of 2.94mm^2^. The bar graph shows the fold increase in both the neoplastic cells (GFPT2-positive) and macrophages and giant cells (PU.1-positive). The three patients all show a strong increase in neoplastic cells and the fraction of macrophages per field remains relatively stable. (e) IHC for GFPT2, representative image from 5 patients that did not receive CSF1R inhibitor therapy (left) and from a patient that was treated with Cabiralizumab (right) shows an increase in GFPT2-positive cells per microscopic field. (f) The number of GFPT2-positive cells from 30 40x fields from 5 untreated patients with a total area of 2.94mm^2^ was compared to 9 fields with a total area of 0.88mm^2^ from a patient after Cabiralizumab treatment. A significant (p<0.05) increase in GFPT2-positive cells was found. (g) Schematic overview showing the decrease in tumor volume due to CSF1R inhibitor treatment that affects macrophages and giant cells but leaves the neoplastic cells unaffected.

A fourth patient was treated with Cabiralizumab, a humanized monoclonal antibody directed against CSF1R. No pretreatment biopsy was available for this patient but a surgical specimen obtained 25 months after cessation of therapy showed a marked increase in the percentage of GFPT2 positive cells that are the neoplastic cells (fig 5e, right panel) when compared to a representative TGCT specimens from untreated patients (fig 5e, left panel). Image analysis of 9 microscopic fields that combined covered an area of 0.88mm^2^ of the first patient’s specimen compared to 30 fields (2.94mm^2^) obtained from 5 different untreated specimens confirmed a marked increase in the percentage of GFPT2-positive cells (fig 5f). Taken together, the findings in the 4 patients suggest that CSF1R inhibitors target the macrophages and giant cells that leads to a decrease in tumor volume. The neoplastic cells are not targeted by the CSF1R inhibitor treatment but are concentrated in a smaller area (fig 5g).

### Molecular support for osteoclast differentiation in TGCT giant cells

The giant cell population in TGCT is often referred to as “osteoclast-like”, a description that is primarily based on their morphology. To further investigate the giant cells in TGCT, we compared their phenotype to giant cell scRNA data that we generated from two Giant Cell Tumors of Bone (GCTB), a tumor with significant osteolytic activity. Like TGCT, GCTB is a neoplasm where a subpopulation of neoplastic cells is associated with bystander cells consisting of macrophages and giant cells. Combined scRNA sequencing data from two GCTB cases yielded a total of 7,358 cells for analysis. The *H3F3A* mutation (G34W) that is pathognomonic for the neoplastic cells was identified in 30 cells (sup fig 6a) and was used to identify the cluster containing the neoplastic cells. The presence of the *H3F3A* mutation corresponded with expression of *TP63* (sup fig 6b) (19) further confirming the identity of the neoplastic cell cluster. The RANK receptor (*TNFRSF11A*) was highly expressed in the osteoclasts and macrophages (sup fig 6d) as described (20, 21). Canonical marker genes were used to identify the remaining clusters (fig 6a). A cell cycle scoring analysis performed with Seurat was used to identify a clusters of proliferating tumor cells and macrophages (sup fig 6c). *CSF1R* was upregulated in the macrophages, giant cells and proliferating macrophages. Moreover, *CSF1* was highly expressed by the neoplastic cell population in GCTB (sup fig 6d), suggesting that CSF1-CSF1R signaling could be an important pathway in GCTB.

**FIG 6.**
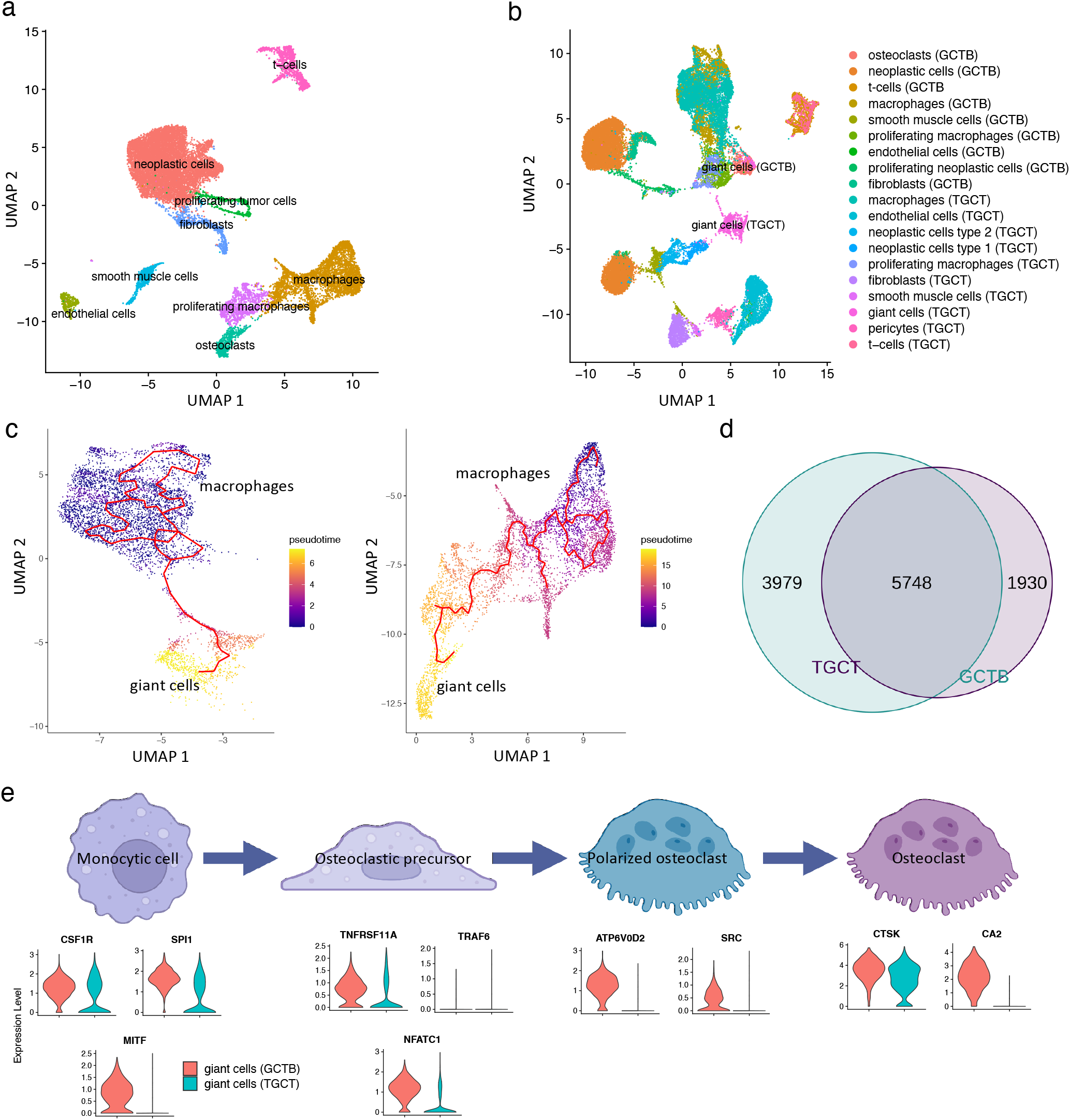
Giant cell development in TGCT is similar to that seen in GCTB. (a) UMAP of scRNA-seq data from 2 GCTB specimens. Clusters were named according to canonical markers, the presence of the *H3F3A* mutation in the neoplastic cells and expression of *TP63* and s scores. (b) UMAP cluster of merged scRNA-seq data from 2 GCTB and 3 TGCT. The giant cells populations cluster near each other but do not overlap. (c) Trajectory analysis from macrophages to giant cells (left, TGCT) and macrophages to giant cells (right, GCTB). (d) The development trajectories in TGCT and GCTB show a significant overlap of 5,748 genes (49%). (e) RNA expression of canonical markers for different development stages of macrophages to osteoclasts are shown for the giant cell clusters in GCTB and TGCT. Giant cells in GCTB are positive for all reported markers except *TRAF6*, while giant cells in TGCT are positive for *CSF1R, SPI1, TNFRSF11A* and *CTSK*.

Integrating the GCTB and TGCT scRNA data and performing a UMAP cluster analysis shows that the giant cell clusters from the two tumor types do not overlap (fig 6b), indicating significant differences between the giant cells from both tumor types. The giant cell clusters from the GCTB and TGCT scRNA samples shared high expression of *SPP1, MMP9* and *CTSK* (sup fig 7a), and IHC for SPP1, MMP9 and CTSK confirmed that all three genes are highly expressed in the giant cell populations of GCTB and TGCT (sup fig 7b).

While the giant cells from both tumor showed no overlap in UMAP cluster analysis, a trajectory analysis identified development trajectories from macrophages to giant cells in both TGCT and GCTB (fig 6c). There is a significant overlap between the genes that change over pseudotime in TGCT and GCTB (fig 6d). These findings suggest that macrophages can develop to giant cells in both TGCT and GCTB through similar developmental mechanisms. Genes associated with osteoclast differentiation were investigated in both GCTB and TGCT giant cells. The giant cells of GCTB were positive for all but one (TRAF6) of the canonical markers that have been described to be associated with osteoclast differentiation (22). The giant cells of TGCT were positive for a smaller but significant subset of these markers with elevated expression of *CSF1R, SPI1, TNFRSF11A, NFATC1* and *CTSK* (fig 6e). IHC further confirmed CTSK expression in the giant cells of GCTB and TGCT (sup fig 7d). CTSK is a known canonical marker gene for osteoclasts (23) and its high expression in TGCT giant cells could explain the bone destruction that is observed in some patients.

## Discussion

TGCT is a rare soft tissue tumor characterized by the presence of a small population of neoplastic cells that harbor a translocation in the *CSF1* gene and that express high levels of *CSF1* mRNA. These cells attract and stimulate a large population of bystander macrophages (3). A number of studies have shown that these tumors react to inhibition of the CSF1 pathway (6–13, 24). To better understand the “landscaping effect” caused by the neoplastic cells on the tumor micro environment we used scRNA-seq and long read RNA-seq. Our data show that there is no CSF1R expression in the neoplastic cells and suggest that the CSF1R inhibitor treatment does not target the neoplastic cells but rather the bystander monocytes. Analysis of patient material from before and after CSF1R inhibitor treatment showed features consistent with a lack of response to CSF1 inhibitors in the neoplastic cells.

Using long read RNA-seq from 3 TCGT cases we found 3 different translocations involving CSF1. We mapped the scRNA-seq data to the discovered translocations and found 79 cells that contained the translocations that were distributed in two clusters. The two neoplastic clusters contained 1160 cells. The relatively low rate of detection of the translocation sequences in scRNA seq data can be explained by the fact that the scRNA-seq libraries are generated by amplification of a variable length mRNA fragments at the 3’ end of genes that may not reach the gene fusion sites in all mRNA strands analyzed. Nevertheless, all cells identified by translocation-specific sequences located to the two cell clusters and thus provided a reliable indicator of the neoplastic nature of the majority of the cells in these clusters. However, we cannot fully exclude that a small subset of the cells in these clusters that could be normal synoviocytes, given the high levels of similarity between these cells. The identity of the neoplastic cells in the two clusters was further confirmed by the high levels of *CSF1* mRNA in the majority of cells. Canonical markers assigned differentiation states to the remaining clusters. A cell cycle state analysis with Seurat was used to identify proliferating cells and showed that the proliferating cell component consisted almost entirely of macrophages, consistent with the known chemoattractant and proliferative effect of CSF1 on these cells.

Analysis of highly expressed genes in neoplastic cells revealed GFPT2 as a marker with a higher specificity for these cells than clusterin. GFPT2 is the rate limiting enzyme in the hexosamine biosynthesis pathway (25). It is expressed in tumor associated fibroblasts in lung cancer (26) and is associated with poor clinical outcome in uterine leiomyosarcoma (27). We showed that GFPT2 co-localized with *CSF1* expression, indicating the presence of the CSF1 fusion (3). The vast majority of the cells in the neoplastic cell cluster express high levels of GFPT2 that is known to lead to activation of the HIPPO signaling pathway (18). The GFPT2-positive neoplastic cells in TGCT show nuclear localization of YAP1 and two downstream genes, indicating activation of the HIPPO signaling pathway. This finding suggests the possibility that the neoplastic cells in TGCT could be directly targeted with HIPPO pathway inhibitors such as Verteporfin (28).

We used immunohistochemistry for GFPT2 to highlight the neoplastic cell in FFPE material and showed that the neoplastic cells have a marked “dendrite-like” morphology. A dendritic appearance of a subset of cells in TGCT was previously reported for clusterin-positive cells (15) and other markers but the dendritic appearance is more pronounced by staining for GFPT2. Previous studies of the *CSF1* fusion pointed to the loss of the 3’-UTR as a possible mechanism for CSF1 upregulation (5, 6). The 3’-UTR contains a negative regulatory sequence that would be lost in translocations involving *CSF1* (5, 6), leading to high levels of expression. The dendritic morphology might facilitate cell-cell integrations and we hypothesized that this could involve a membrane-bound form of CSF1 on the neoplastic cells and CSF1R on the bystander macrophages. CSF1 contains a cleavage site in exon 6 followed by a transmembrane domain. This cleavage site is used to release CSF1 from the cell surface, without this cleavage site CSF1 is membrane bound (29). In our 3 cases, long read RNA-seq elucidated the exact translocation break points and translocation partners for the *CSF1* gene. In two cases the transmembrane domain and cleavage sites were lost but the translocation partner donated either a transmembrane domain (in case of PDPN) or a fibronectin domain that likely binds to the cell surface (in case of FN1) (30). The third TGCT case had a translocation involving the 3’-UTR of CSF1. In this case 20% of the reads show a splice variant that lacks the exon 6 transmembrane domain (data not shown). The sequences from CSF1 translocations reported by Tap et al. (6) also point to the loss of the CSF1 transmembrane domain and cleavage site but also the potential gain of other transmembrane domains from the translocation partners, for example in the translocations involving CD99 and CD101 as fusion partners. The presence of the membrane bound CSF1 could explain the advantage of a dendritic morphology in the neoplastic cells to facilitate interaction with the macrophages and further studies are required to address this issue.

To further study the relations between the different cell types that comprise TGCT, receptor-ligand interaction analysis was performed for CSF1 and PDGF pathways and this suggested a strong interaction between CSF1 produced by neoplastic cells and the CSF1R present on macrophages and giant cells. No interaction between CSF1 and CSF1R on the neoplastic cells was found and indeed no CSF1R expression was found on the neoplastic cells, providing no support for an autocrine loop using the CSF1 pathway. Clinical trials showed that CSF1R inhibitors such as pexidartinib are effective targeted therapies for TGCT and show a reduction in tumor mass measured by MRI according to the Response Evaluation Criteria in Solid Tumors (RECIST) or TVS criteria (6, 12, 14). The lack of CSF1R on the neoplastic cells suggests that the CSF1R inhibitors target the bystander macrophages and giant cells but not the neoplastic cells. Patient samples from before and after CSF1R inhibitor treatment were studied by identifying the neoplastic cells, macrophages and giant cells on FFPE sections. We showed an increase in the fraction of neoplastic cells while the fraction of macrophages remained stable measured in comparably sized histological sections. The radiologically measured TVS decreased (2/3) or remained stable (1/3). Based on our data it seems likely that the bystander macrophages and giant cells are targeted by the CSF1R inhibitor treatment causing a significant shrinkage of the tumor and concentrating the neoplastic cell population within a smaller volume. The increase in fraction of neoplastic cells could also be caused by proliferation but in untreated samples we were unable to find expression of proliferation markers in the neoplastic cells. It is not possible to quantify the absolute number of neoplastic cells in the tumor before and after treatment as we are limited to histological sections. The analysis of the few cases for which samples from before and after kinase inhibitor treatment was available is further complicated by the fact that there was significant variation in the time between cessation of therapy and the resection of the post-treatment sample. Another variable is the different response rates to CSF1 pathway inhibitors seen in patients. It is not unlikely that after stopping CSF1R inhibitor treatment the neoplastic cells would attract new macrophage and giant cell populations and over time would regain the volume. Tap et al describe a future trial that will look at the long term effects of discontinuation and retreatment with CSF1R inhibitor treatment (6) that may address this issue.

In addition to macrophages, TGCT also contain multinucleated giant cells. These cells have not been well characterized but are referred to as “osteoclast-like” based on their morphology. To our surprise the cell size restrictions for the 10X Genomics Chromium microfluidics did not exclude giant cells from our single cell sequencing library and we were able to identify a cluster of multinucleated giant cells in our scRNA-seq data. Studying H&E-stained smears that were prepared after tumor dissociation and size filtering for cells that would pass through the 10X Genomics Chromium microfluidics showed the presence of the giant cells on these slides (sup fig 5b bottom panel), showing that the giant cells could pass through the filter. In our scRNA data we found further convincing support for the successful analysis of giant cells. First, levels of MMP9, SPP1 and C1QA mRNA correspond to IHC findings on giant cells in TGCT. Second, in the Giant Cell Tumor of Bone (GCTB) specimens we could identify the giant cells by their expression of TNFSF11, a characteristic marker for these cells in this tumor. Third, while the giant cells in TGCT and GCTB appear quite large in histologic images, the giant cells in GCTB have been measured as having size range of 10-300um (31). Finally, other groups were also able to study giant cells using single cell sequencing, such as the single cell study of GCTB (32, 33). The presence of giant cells in the scRNA data allowed to compare them to those found in Giant Cell Tumor of Bone (GCTB) that has several similarities with TGCT. Both are characterized by a population of neoplastic cells driving formation of a tumor consisting mainly of macrophages and giant cells (34). In GCTB destruction of bone is very common but this is usually not a prominent finding in TGCT. Recent single cell studies of GCTB showed how monocytes developed into osteoclasts (32). In TGCT the giant cell population are described as “osteoclast-like giant cells” but it is unknown to what extend the multinucleated giant cells are similar to osteoclasts or to those giant cells found in GCTB. In addition, the possible path for development from TGCT monocytes into giant cells has not been studied. Trajectory analysis showed such a path for development to multinucleated giant cells in TGCT that is similar but not identical to what was recently reported for giant cells in GCTB (32). There were significant differences between the giant cells found in TGCT as compared to GCTB. The giant cells did not cluster together when the data was integrated and there were differences in expression of osteoclast development markers showing that there are significant differences between the two giant cell populations. However, both giant cell clusters from TGCT and GCTB express high levels of CTSK, as seen with scRNA-seq and IHC, which is described to be a specific marker and important functional protein in osteoclast development (23). CTSK is a potential target on the giant cell population for which inhibitors have been developed (35, 36).

In conclusion, we used scRNA-seq to perform a detailed study of the tumor landscape of TGCT. By integrating our data with a published scRNA dataset derived from the synovium of patients with rheumatoid arthritis we found a remarkable overlap between the 2 neoplastic cell clusters in TGCT and the two types of synoviocytes providing proof that the neoplastic cells in TGCT are derived from synoviocytes. We found that GFPT2 is a sensitive and specific biomarker for the neoplastic cells. We show that the neoplastic cells stimulate bystander macrophages and giant cells through CSF1-CSF1R signaling, possibly through direct cell contact rather than secreted CSF1. There is no evidence of an autocrine feedback loop and it is unlikely that the CSF1R inhibitor treatment targets the neoplastic cells, this was further supported by findings in patient material from before and after treatment with CSF1R inhibitors. This observation has significant clinical consequences as the risk for recurrence after cessation of treatment may be high. Very little data on the rate of recurrence is currently known. The giant cell population was characterized and compared to the giant cells in GCTB. Our data further shows that inhibiting the HIPPO signaling pathway could directly target the neoplastic cell population in TGCT and that the giant cell population could respond to CTSK inhibitors.

## Patients and Methods

### Patient material

Tumor material from three TGCT cases (diffuse type) and 2 cases of GCTB were used for scRNA sequencing. Six Stanford TGCT cases were used for validation and three TGCT cases from the Leiden University Medical Center with material from before and after CSF1R inhibitor treatment were included. The study was approved by the Stanford University Institutional Review Board (IRB-58498). Material from the Leiden University Medical Center was derived from the bone and soft tissue tumor biobank and coded according to the ethical guidelines described in “Code for Proper Secondary Use of Human Tissue in The Netherlands” of the Dutch Federation of Medical Scientific Societies as approved by the Medical Ethical Board (B22.010).

### Tumor dissociation and single cell RNA-seq

To dissociate fresh tumor tissue for single cell RNA-sequencing 250mg of tissue was minced and dissociated according to the protocol for the Tumor Dissociation Kit (130-095-929; Miltenyi Biotec) using a GentleMACS Octo Dissociator with heaters (130-095-937; Miltenyi Biotec) using TDK setting 3. The dissociated cell suspension was then applied to a 70um MACS SmartStrainer (130-098-462; Miltenyi Biotec). The quality of the suspension was checked by first generating a H&E slide from a smear of the suspension. Secondly, the sample was analyzed using an automated cell counter (TC10; BioRad). Samples were further processed by the Stanford Functional Genomics Facility. Single cell libraries were sequenced on the NovaSeq6000 platform (Illumina) on a S1 Flow Cell (Illumina).

### Long read RNA-Seq

RNA was isolated from TGCT1-3 using the RNeasy Plus Mini Kit according to the manual (74134; Qiagen). The three samples were pooled into one cDNA library and sequenced on one SMRT cell (PacBio). PacBio Iso-Seq sequencing was performed on the Sequel 2 machine (PacBio).

### Immunohistochemistry and immunofluorescence

For both immunohistochemistry (IHC) and immunofluorescence (IF), 4um paraffin sections were deparaffinized with xylene. Slides were re-hydrated using decreasing concentrations of Ethanol. Antigen retrieval was performed at 121°C with the slides in either EDTA Ph9 or Citrate Ph6. Primary antibodies for GFPT2 (1:500; ph9; ab190966; Abcam), PU.1 (1:50; ph6; 554268; BD Biosciences), CD68 mouse (1:200; ph9; ab955; Abcam), CD68 rabbit (1:200; ph9; 76437S; Cell Signaling), MMP9 (1:1000; ph9; HPA001238; Millipore Sigma), YAP1 (1:1200; ph6; sc-101199; Santa Cruz), desmin (1:40; D33 Dako) and Ki67 (1:50; ph9; Cell Signaling) were used. For IHC immunodetection was completed using the Vectastain ABC kit (Vector Laboratories) and DAB chromogen (Abcam), according to the manufacturer’s specifications. For IF a secondary mouse or rabbit Alexa488, Alexa555 or Alexa657 labeled antibodies were used (ThermoFisher). IHC and IF sections were scanned using the Keyence Fluorescence Microscope (BZ-X; Keyence) with the 40x lens, either using the fluorescent filter cube or the brightfield setting.

### Fluorescent In Situ Hybridization

FISH for CSF1 was performed as previously described (3). BACs (CSF1: 354C7, 19F3 and 96F24) from the Human BAC Library RPCI-11 (BACPAC Resources Center, Children’s Hospital Oakland Research Institute) were used.

### RNA In Situ Hybridization combined with immunohistochemistry

RNA-ISH was combined with IHC using the RNA Protein Co Detection Assay (323180; ACD Bio) combined with RNAscope® 2.5 HD Detection Kit - RED (322360; ACD Bio) according to the provided protocol. GFTP2 antibody was used at a concentration of 1:50 and *CSF1R* and *CSF1* probes were ordered from ACD Bio.

### Bioinformatics

#### Analysis software and visualization

R (v4.1.0) and Python (v3.9.7) were used for bioinformatic analysis. Plots were generated with Seurat, ggplot, Seaborn or Prism (v9.3.1; GraphPad).

#### Single cell RNA-seq data processing and quality control

Raw scRNA-seq data was pre-processed using the Cell Ranger workflow (v6.1.2; 10X Genomics). The raw_feature_bc_matrix object was imported into Seurat (v4.1.0) (37). Low-quality cells were excluded; cells with fewer than 200 features or less than 10% mitochondrial RNA were removed. The data was normalized using the NormalizeData function in Seurat. The single cell expression data was clustered using the UMAP algorithm RunUMAP in Seurat. Seurat was used to identify clusters with a resolution setting of 0.4. The clusters were named by identifying the differentially expressed genes in each cluster using the FindAllMarkers function in Seurat. Only upregulated genes were identified that had at least a 0.25 log2fold change and an adjusted p<0.05. The differentially expressed genes were manually screened for known cell-differentiation in the literature.

#### Combining scRNA-seq datasets

After normalizing the different RNA-seq libraries anchor points were identified using the FindIntegrationAnchors function in Seurat and the data was integrated using the IntegrateData function. Using the RunUMAP and FindClusters functions the new dataset was analyzed for cell clusters.

#### Support Vector Machines classifier

The R package scPred (v1.9.2) (38) was used to create a Support Vector Machine (SVM) to identify synoviocytes subtypes 1 and 2. The SVM classifier was subsequently applied to the neoplastic cell clusters.

#### Trajectory analysis

Monocle3 (v3.1)(39) was used to infer development trajectories between macrophages and osteoclasts in GCTB and macrophages and giant cells in TGCT samples. The relevant cells were subset and re-clustered using the cluster_cells function with a resolution setting of 1e-4. Development paths were identified using the learn_graph function. Using order_cells the pseudotime was inferred.

#### Receptor-ligand interaction analysis

CellphoneDB (v3)(40) was used to identify cell-cell interaction in the scRNA-seq data. The read counts and meta data were exported from the Seurat object. Receptor-ligand interactions were filtered for significant (p<0.05) interactions involving neoplastic cells and other cell types.

#### PacBio Iso-Seq analysis and fusion detection

PacBio Iso-Seq data was aligned to hg38 (UCSC) using Minimap2 (v2.24)(41) with the recommended settings for Iso-Seq data. Squanti3 (v4.3)(42) was used to extract fusion genes from the aligned bam files. Identified fusions were manually screened using IGV (v2.4.1; Broad).

#### Translocation detection in scRNA-seq data

The sequences for the identified fusions identified in TGCT cases 1-3 were used to create a Bowtie 2 (v2.4.5)(43) reference genome. The reads from the scRNA-seq experiments were aligned to the custom fusion references using Bowtie 2. Samtools view (v1.15.1) was used isolate break-point spanning reads and these reads were further validated by hand using the IGV.

#### Immunofluorescence analysis

Scanned images were analyzed using OpenCV-python (v4.5.5.64). To quantify IF signals first an Otsu threshold was used on the DAPI channel to identify the nuclei locations and determine the number of cells. The intensity of staining in the corresponding cytoplasm was determined by dilating the nucleus location and subtracting the area of the nucleus. IF signal intensity was determined for the different IF markers in the cytoplasm or nucleus masks applied to the corresponding IF stain imaging data.

## Figure legends

**SUP FIG 1.**
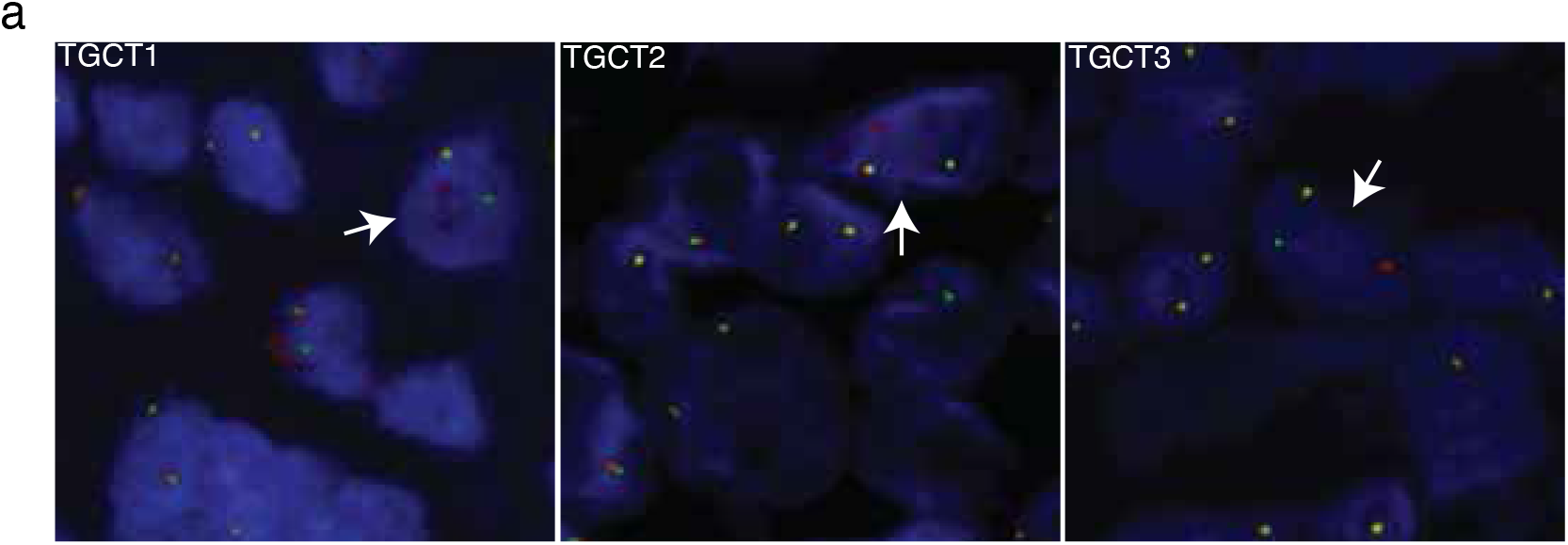
Fluorescence In Situ Hybridization confirms the presence of the CSF1 fusion in a subset of cells. Arrows indicate cells with break-apart green and red signal (flanking *CSF1*), while the probes for the other chromosome co-localize.

**SUP FIG 2.**
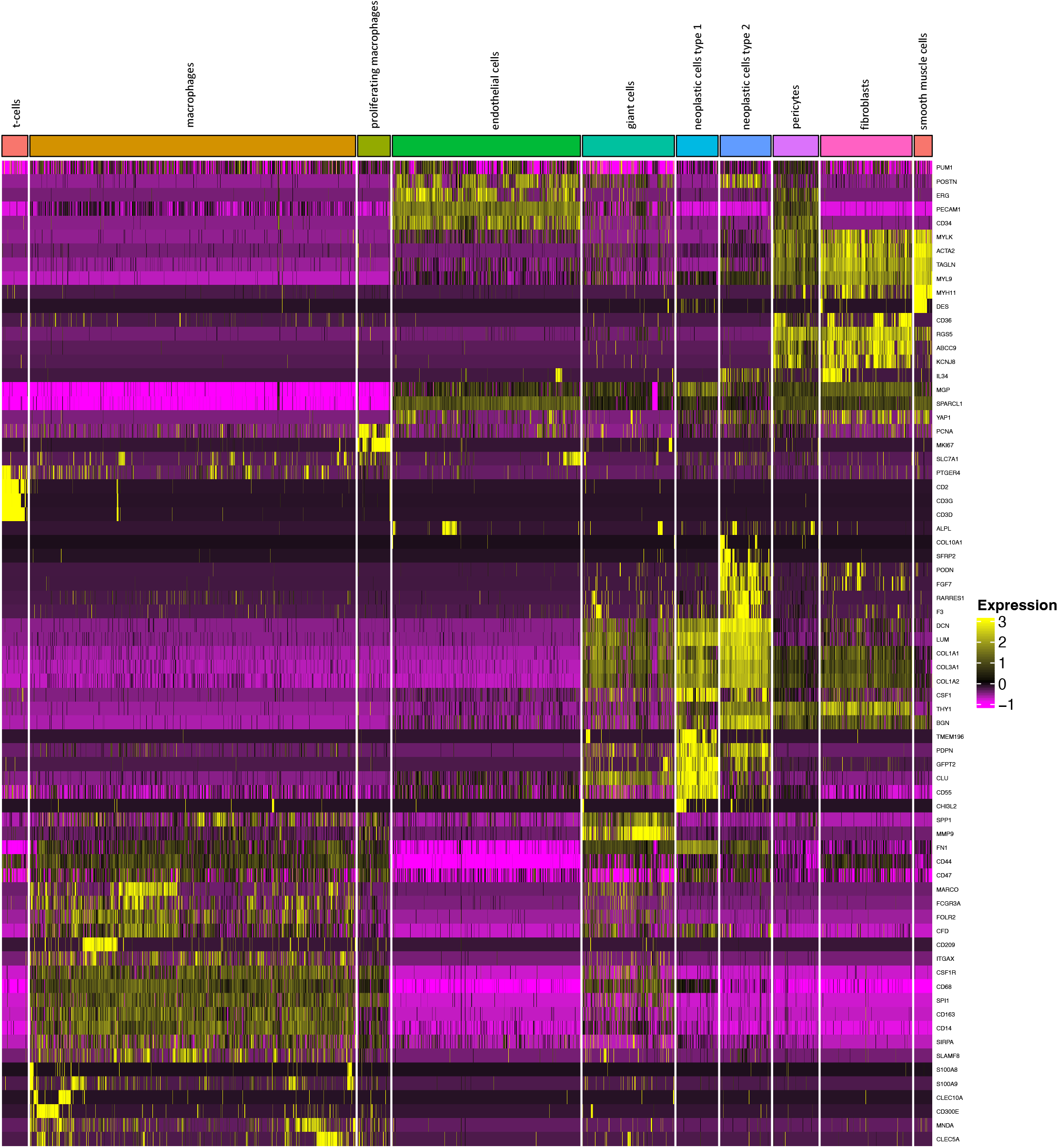
The canonical markers used to identify the cell types corresponding to the different clusters in TGCT.

**SUP FIG 3.**
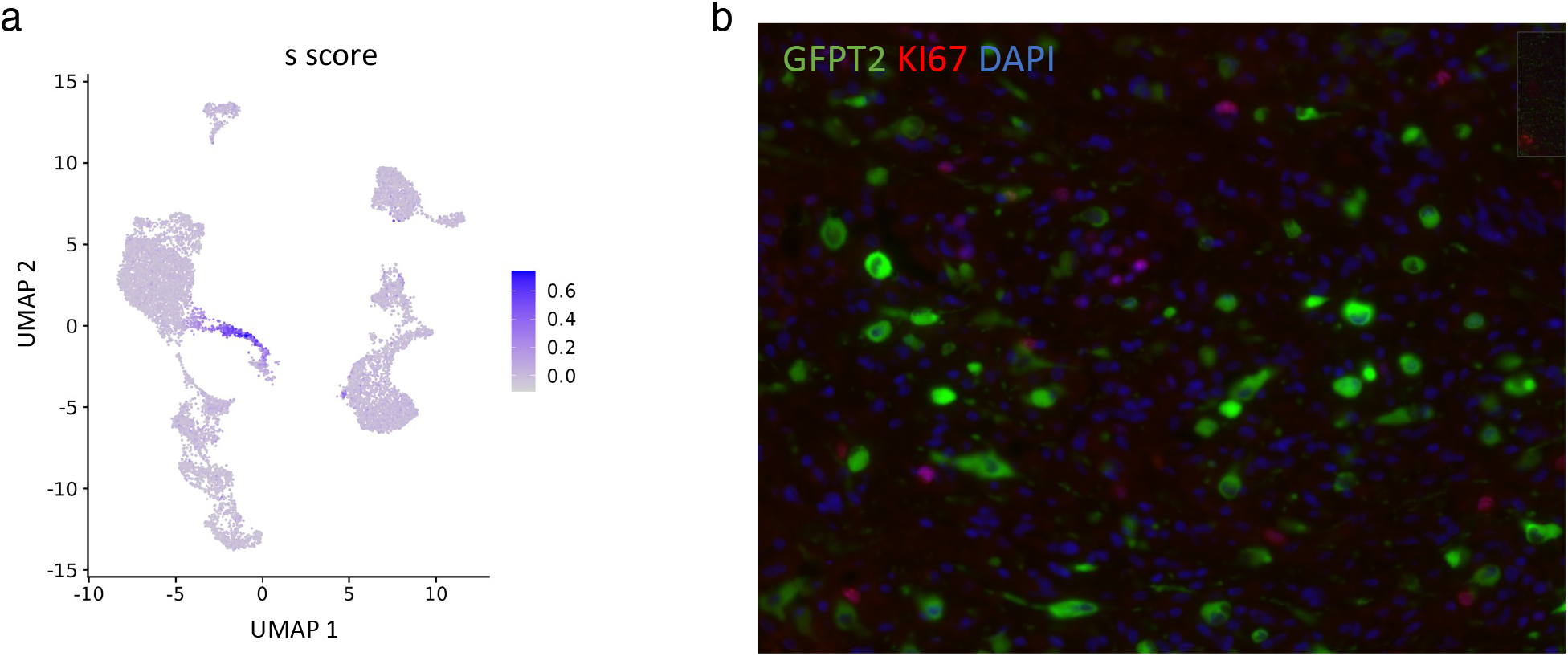
In TGCT proliferation is seen in a subset of the macrophages but not in the neoplastic cells. (a) Scatterplot of the scRNA-seq data from TGCTs shows high s scores in a subset of the macrophages, the neoplastic cells are negative. (b) Immunofluorescence stain for KI67 and GFPT2 shows that GFPT2 + cells (neoplastic cells) are negative for KI67 which is positive in some of the surrounding cells.

**SUP FIG 4.**
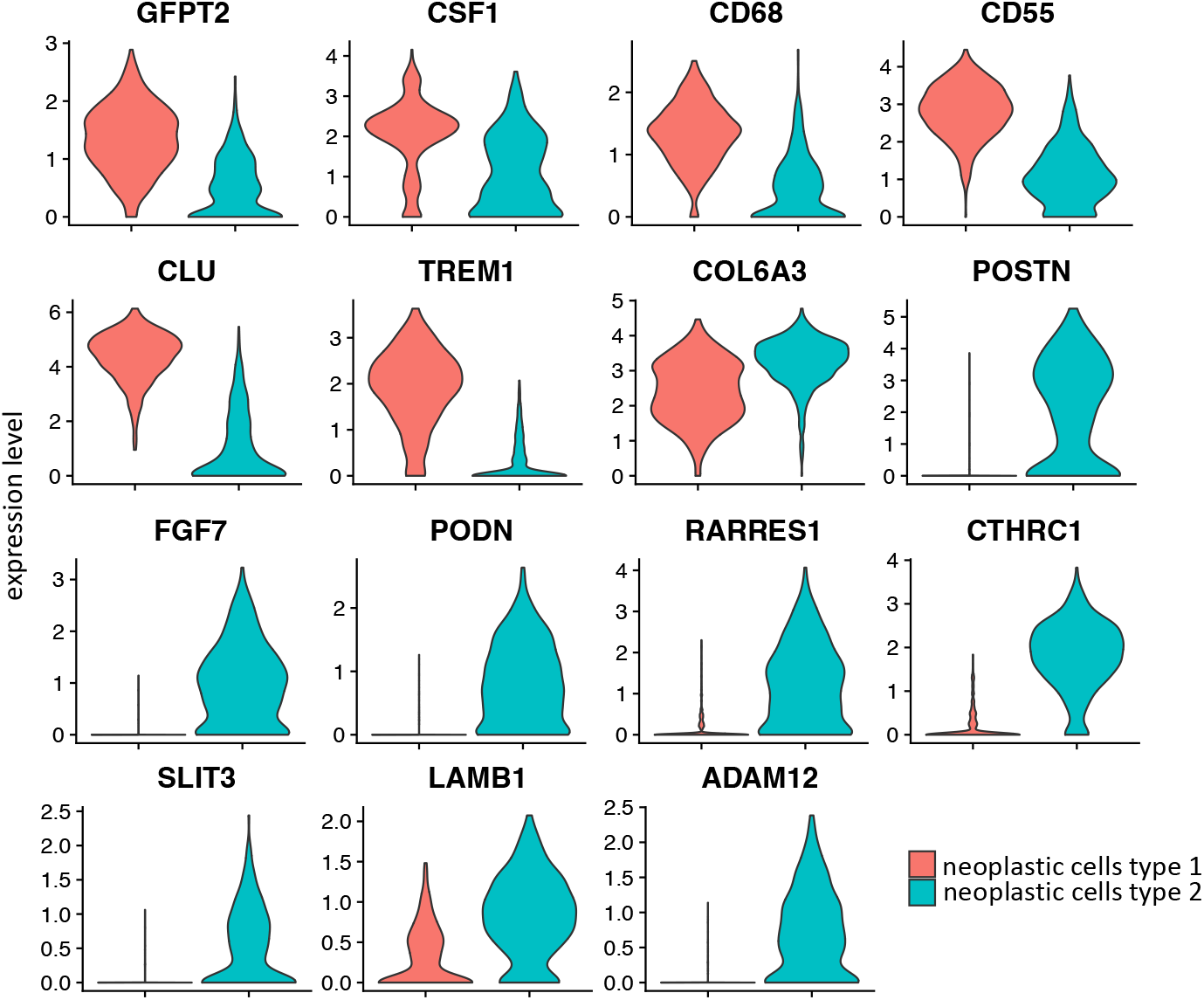
Differentially expressed genes between the two neoplastic cell clusters in TGCT.

**SUP FIG 5.**
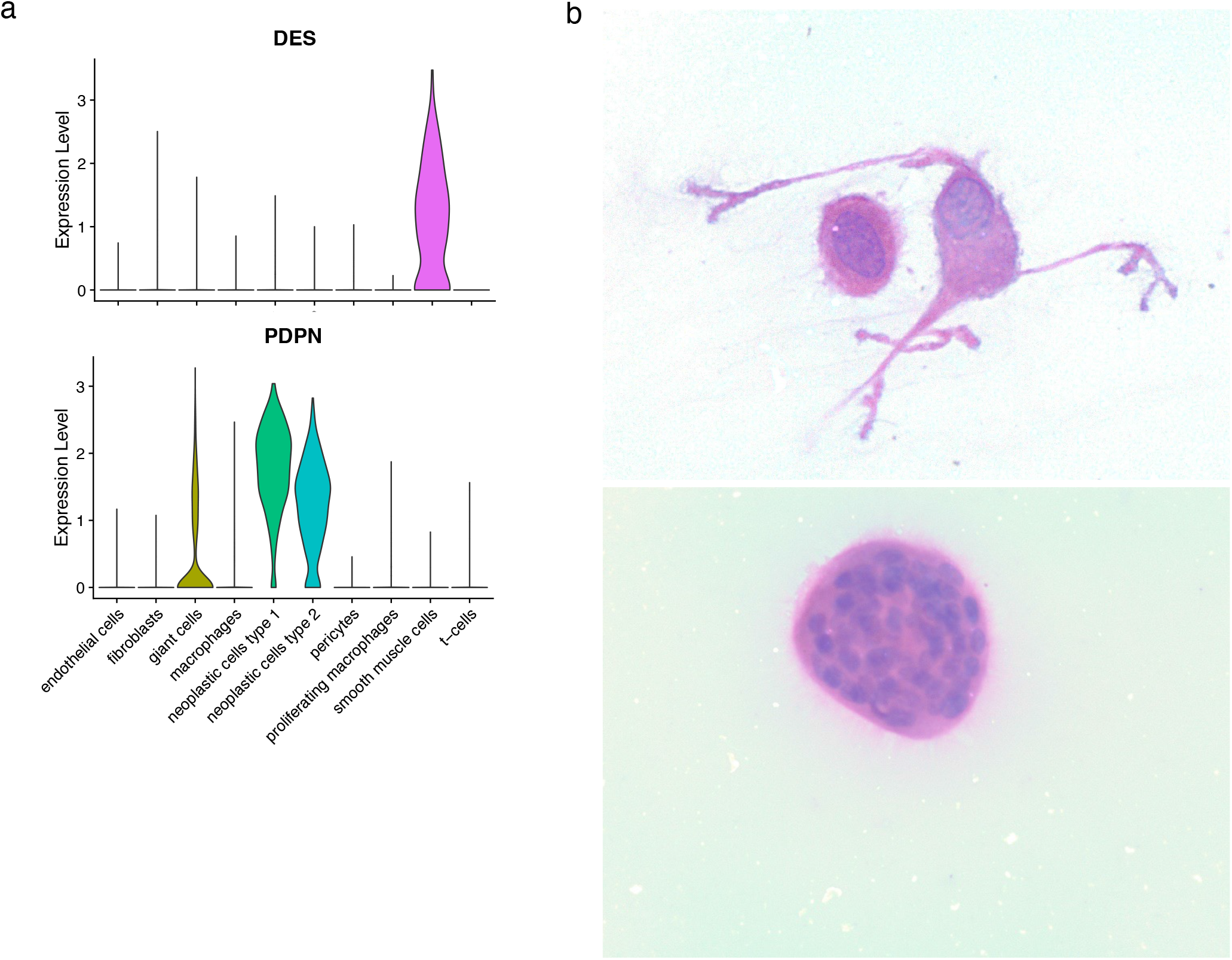
(a) Expression of *DES* and *PDPN* in the different TGCT clusters. (b) H&E-stained smear of a single cell preparation of TGCT after straining shows cells with a dendritic morphology (top) and a multinucleated giant cell (bottom).

**SUP FIG 6.**
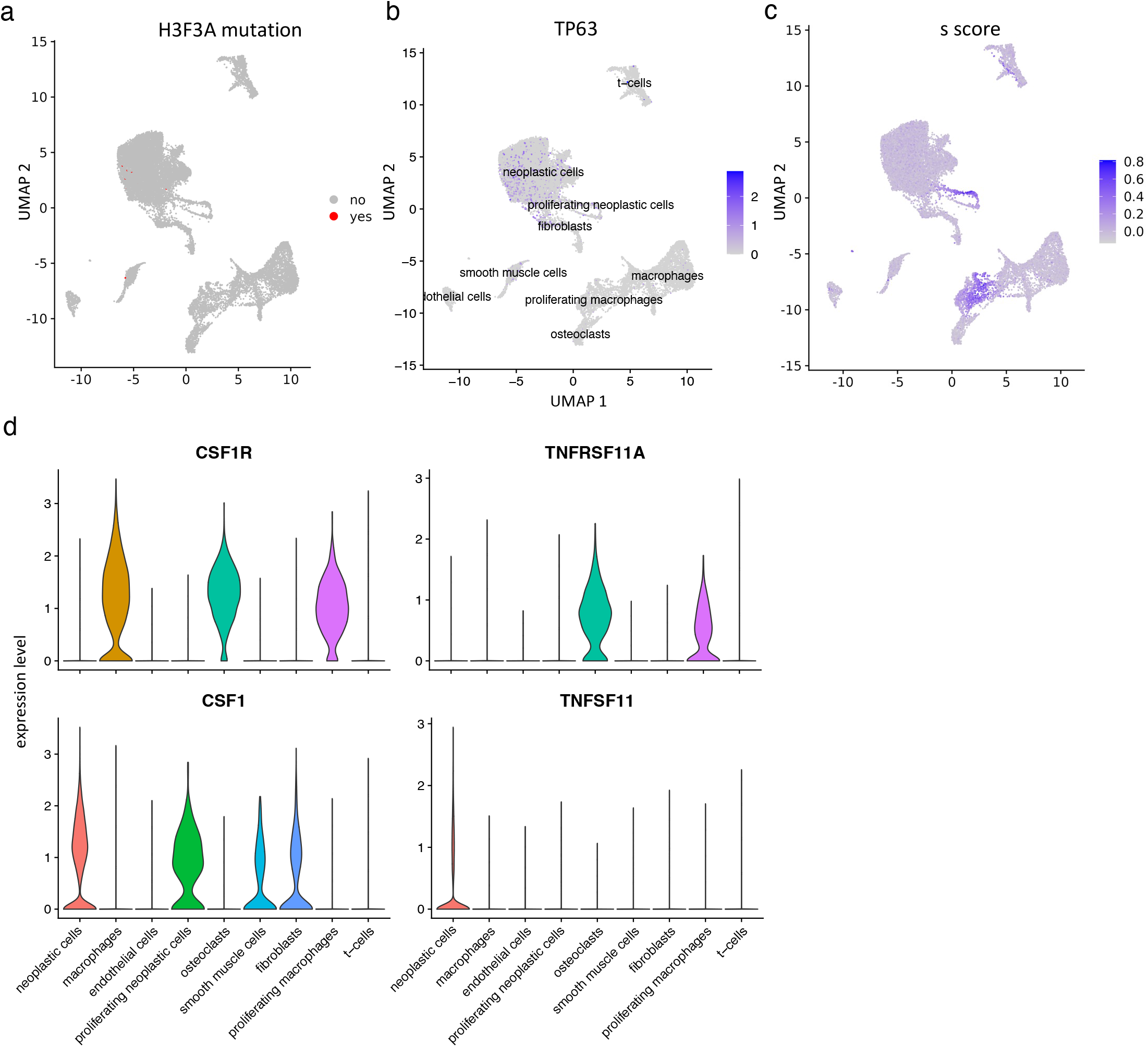
The different cell types present in GCTB were identified using canonical markers, the presence of *H3F3A* mutations, expression of *TP63* and s scores. Neoplastic cells interacted with the macrophages and giant cells through RANK (*TNFSF11*) and *CSF1* signaling. (a) Cells with *H3F3A* mutation were identified in the scRNA-seq data (red). (b) Scatter plot showing the presence of *TP63* positive cells. (c) Scatter plot showing a high s score in a subset of macrophages and neoplastic cells. (d) Gene expression genes involved in RANK and CSF1 signaling.

**SUP FIG 7.**
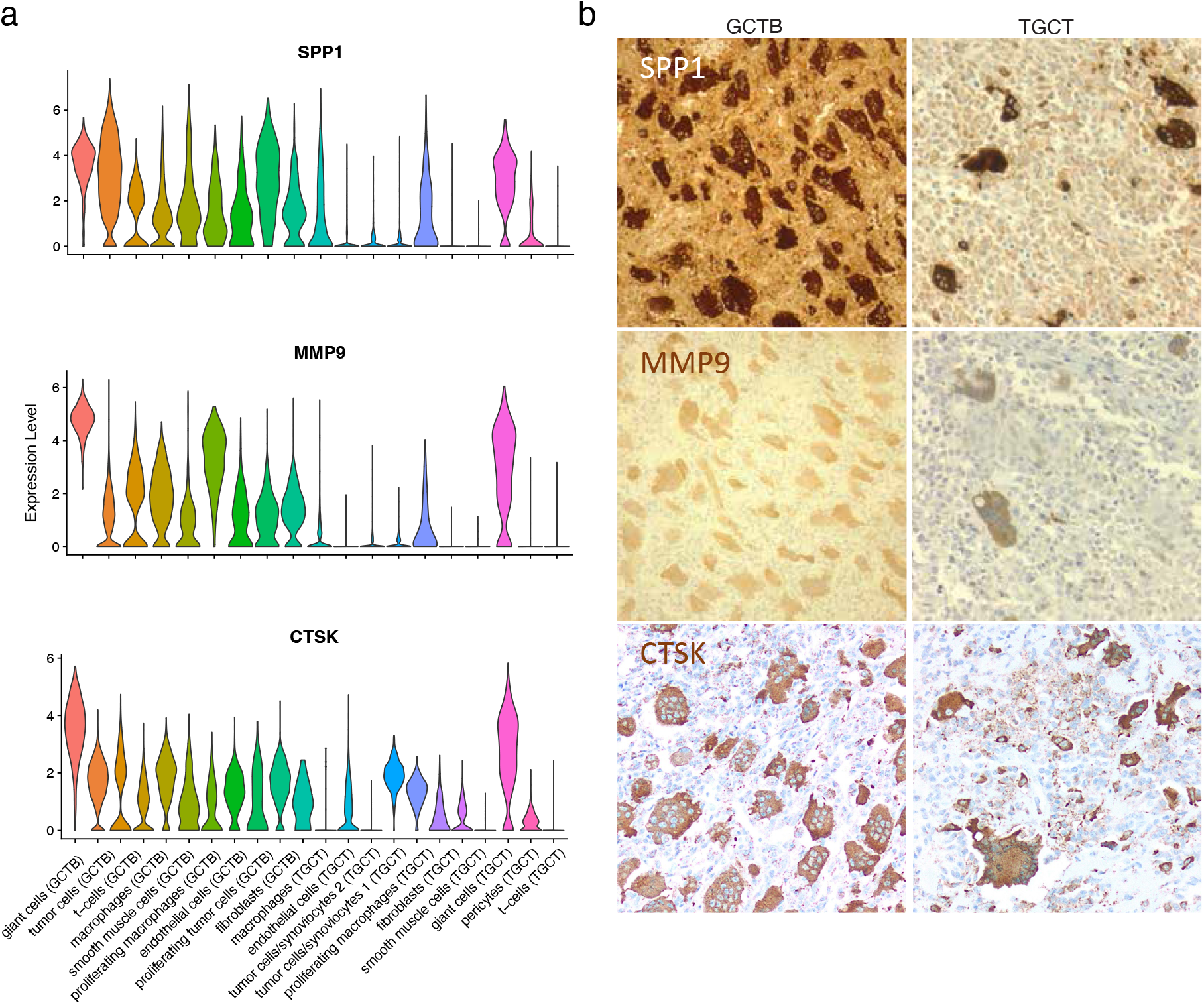
Giant cells in GCTB and TGCT show expression of similar marker genes for which expression was confirmed on protein level. (a) Expression of *SPP1, MMP9* and *CTSK* in GCTB and TGCT. (b) IHC for SPP1, MMP9 and CTSK.

**SUP TABLE S1.**
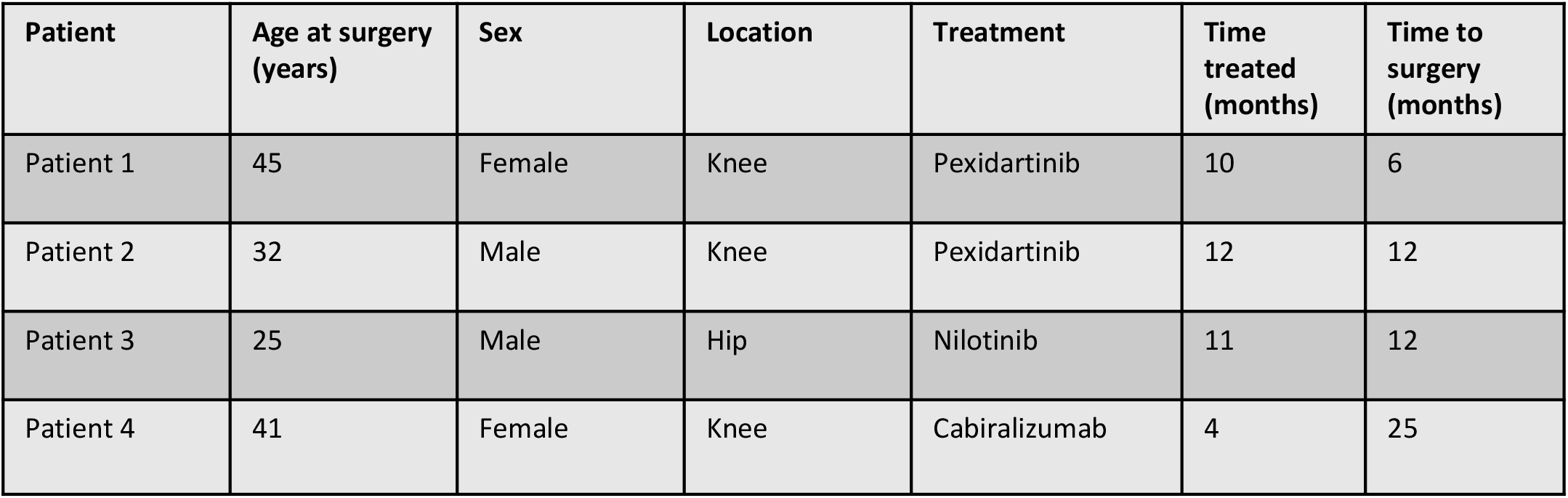
Clinical information for the patients that received CSF1R inhibitor treatments.

## Acknowledgements

The sequencing data was generated with instrumentation purchased with NIH funds: S10OD025212 and 1S10OD021763. This research was supported by an unrestricted grant of Stichting Hanarth Fonds, The Netherlands to DVIJ. This research was supported by the Virginia and D.K. Ludwig Fund for Cancer Research and the Taube Family Foundation. Figures 1b, 5g and 6e created with BioRender.com.

